# Cardiomyocytes upregulated PD-L1 expression to alleviate cardiac injury induced by irradiation combined with anti-PD-1 antibody: an in vitro and in vivo study

**DOI:** 10.64898/2026.09.14.751375

**Authors:** Ye Zhou, Yajing Wu, Na Zhang, Shuo Wang, Xueshuai Ye, Jingtao Ma, Qingxia Li, Jun Wang

## Abstract

**Objective:** Preclinical studies suggest that irradiation combined with anti-PD-1 antibody (iRT) exacerbates cardiac injury in mice, partly through CD8⁺ T-cell-mediated mechanisms. We investigated radiation-induced PD-L1 expression in cardiomyocytes and explored its potential role in iRT-associated cardiac injury.

**Methods:** AC16 cells were exposed to different doses of irradiation. PD-L1 expression was assessed at different time points by flow cytometry, qRT-PCR, western blotting, and immunofluorescence. Cell viability, apoptosis, and cytokine levels were evaluated. PD-L1 was knocked down using siRNA to assess its function. PBMCs or CD8⁺ T cells were co-cultured with irradiated or non-irradiated AC16 cells at different E:T ratios, with or without PD-1 blockade. A C57BL/6 mouse model of iRT-induced myocardial injury was established, and myocardial PD-L1 expression and T-cell infiltration were evaluated.

**Results:** Irradiation upregulated PD-L1 expression in AC16 cells, with increased membrane and cytoplasmic signals at 48 h and prominent nuclear-associated fluorescence at 72 h. Irradiation increased IL-6, MCP-1, CCL5, and CXCL10 release, decreased IL-10, reduced cell viability, and increased apoptosis. PD-L1 knockdown further increased inflammatory cytokine and chemokine release but did not significantly alter apoptosis or further reduce viability in irradiated AC16 cells. In the presence of a high density of activated CD8⁺ T cells, disruption of PD-1/PD-L1 signaling was associated with a further reduction in AC16 cell viability. In vivo, myocardial PD-L1 expression was increased, whereas CD8⁺ T-cell infiltration remained limited and focal.

**Conclusion:** PD-L1 upregulation in cardiomyocytes may exert an adaptive protective and immunoregulatory role during iRT-associated injury, potentially by limiting inflammatory mediator release and modulating susceptibility to activated CD8⁺ T-cell-mediated injury. CD8⁺ T-cell density and activation are the key issues.

## Introduction

Radiotherapy for thoracic tumors should minimize cardiac exposure, and radiation-induced heart disease (RIHD) has become an important concern during treatment planning. In RTOG 0617, higher cardiac radiation exposure was associated with worse overall survival ^[1]^. In a separate pooled analysis of six prospective dose-escalation trials in stage III non-small-cell lung cancer (NSCLC), Wang et al. reported symptomatic cardiac events in 26 of 112 evaluable patients (23%), with estimated event rates of 14% at 2 years and 32% at 4 years ^[2]^.

Immune checkpoint inhibitor (ICI)-related cardiac toxicity has garnered significant attention because of its high case fatality ^[3–5]^. Among 36,848 immunotherapy toxicity reports submitted to the U.S. Food and Drug Administration Adverse Event Reporting System (FAERS) from 2017 to 2018, 2,316 involved cardiovascular toxicity and 816 had a fatal outcome, corresponding to a fatal-outcome proportion of 35.2% among reported cardiovascular toxicities; myocarditis had a reported fatality of approximately 50% ^[5]^. Pre-existing cardiovascular conditions have also been associated with increased reporting of ICI-related myocarditis in pharmacovigilance data ^[6]^.

When radiotherapy and immunotherapy are combined, it remains unclear whether cardiac toxicity is increased. Myers et al. reported that an anti-PD-1 antibody combined with thoracic irradiation (iRT) resulted in worse survival in mice, associated with increased T-cell infiltration in cardiac and pulmonary tissues ^[7]^. Bai et al. similarly reported exacerbated myocardial injury, fibrosis, and cardiomyocyte apoptosis in a mouse iRT model ^[8]^. Clinical evidence has been less consistent. A pooled analysis of prospective ICI trials in the FDA database found no meaningful increase in serious adverse events among patients who received radiotherapy within 90 days before ICI treatment ^[9]^, and a small retrospective lung-cancer study did not identify a significant increase in cardiac events with thoracic radiotherapy plus ICI ^[10]^. In PACIFIC, durvalumab after concurrent chemoradiotherapy improved 3-year overall survival to 57.0% versus 43.5% with placebo ^[11]^; deaths attributed to adverse events occurred in 4.4% and 6.4% of patients, respectively ^[12]^.

Discrepancies between preclinical animal models and clinical findings highlight the need for further investigation. Du et al. reported a 30% acute mortality within 2 weeks following combined cardiac irradiation and anti-PD-1 treatment, compared with 0% mortality in cardiac irradiation plus immunoglobulin G-treated mice ^[13]^. This acute toxicity was mediated by CD8⁺ T cells, as depletion of these cells using anti-CD8 antibodies reversed the mortality. A clinical case report described a patient with multiple myeloma who developed fatal myocarditis after combined radiotherapy and ICI therapy, coinciding with an increased population of peripheral blood PD-1^hi^Ki-67^hi^ CD8⁺ T cells ^[14]^. These observations highlight the potential role of CD8⁺ T cells in iRT-related cardiac toxicity.

Interestingly, PD-L1 can be expressed in human cardiomyocytes and is inducible in murine cardiac myocytes during inflammatory injury ^[15,16]^. The PD-1/PD-L1 pathway has been shown to restrain cardiac inflammation in autoimmune myocarditis ^[17]^, viral myocarditis ^[16]^, and CD8⁺ T-cell-mediated myocardial injury ^[18]^. PD-L1-deficient MRL mice develop autoimmune myocarditis ^[17]^, whereas PD-1-deficient MRL mice develop fatal myocarditis with massive CD4+ and CD8⁺ T-cell infiltration ^[19]^. These studies suggest that the PD-1/PD-L1 axis may mitigate myocardial injury by restraining excessive immune responses ^[19–21]^. However, the specific role and mechanisms of the PD-1/PD-L1 pathway in myocardial injury induced by iRT remain incompletely understood.

Previous studies have reported that radiation can contribute to myocardial injury by inducing elevated levels of inflammatory factors and activating immune cells surrounding cardiomyocytes^[22]^. Furthermore, in chronic Chagas cardiomyopathy, a severe inflammatory cardiomyopathy, upregulation of proinflammatory chemokines such as CXCL10, CCL5, and MCP-1 have been contributes to myocardial autoimmune loops ^[23]^. Despite these observations, how irradiation regulates PD-L1 expression in cardiomyocytes and how this response interacts with inflammatory signaling and CD8⁺ T-cell-mediated injury remain incompletely understood.

To address these questions, we investigated radiation-induced changes in PD-L1 expression and subcellular distribution in human cardiomyocytes and examined the effects of PD-L1 disruption on inflammatory mediator release and cardiomyocyte viability. We further evaluated how the density and activation status of CD8⁺ T cells, together with disruption of PD-1/PD-L1 signaling, influenced cardiomyocyte injury in a co-culture system. Finally, myocardial PD-L1 expression and T-cell infiltration were assessed in a mouse model of localized cardiac irradiation with or without PD-1 blockade. Through these complementary experiments, we aimed to characterize the potential adaptive immunoregulatory role of cardiomyocyte PD-L1 during radiation-associated inflammatory stress.

## Materials and Methods

### Animals and Cell Lines

AC16 cells were procured from Millipore and cultured in DMEM supplemented with 10% fetal bovine serum (FBS), 1% penicillin-streptomycin, and 2 mM glutamine. Male C57BL/6 mice (8 weeks old, 20 g) were obtained from the Fourth Hospital of Hebei Medical University and housed under standardized laboratory conditions. All experimental procedures involving animals were reviewed and approved by the Animal Ethics Committee (No. 2022035).

### Anti-PD-1 Antibodies

Camrelizumab, a humanized anti-PD-1 monoclonal antibody, was obtained from Hengrui Pharmaceuticals (Jiangsu, China) and used for PD-1 blockade in human T-cell co-culture experiments. A mouse-reactive anti-PD-1 monoclonal antibody (HRP00232-025) was provided as a gift by Hengrui Pharmaceuticals (Jiangsu, China) and was used for the in vivo experiments.

### In Vitro Model of Radiation-Induced Human Cardiomyocyte Injury

AC16 cells were cultured in high-glucose DMEM supplemented with 10% FBS at 37°C in a humidified atmosphere containing 5% CO₂, with media replacement every 48 hours. Cells were trypsinized, centrifuged, resuspended, and seeded into six-well plates at a density of 2.5 × 10⁵ cells/mL. Following 6 hours of incubation to facilitate cell attachment, the cultures were subjected to irradiation doses of 6 Gy or 10 Gy using a Varian linear accelerator to establish an in vitro model of radiation-induced cardiomyocyte injury.

### In Vivo Model of iRT-Induced Myocardial Injury in Mice

Twenty male C57BL/6 mice (8 weeks old, 20 g) were randomly allocated into four experimental groups: Group A (control, no treatment), Group B (treated with PD-1 monoclonal antibody), Group C (subjected to 20 Gy cardiac irradiation with saline injection), and Group D (treated with anti-PD-1 antibody in combination with 20 Gy cardiac irradiation). Mice were anesthetized and exposed to 20 Gy of localized cardiac irradiation using a Varian Clinac 23Ex linear accelerator. Mice in Groups B and D received intraperitoneal injections of anti-PD-1 monoclonal antibody (10 mg/kg) every other day for a total of 14 doses. On day 28 post-irradiation, all animals were euthanized, and cardiac tissues were harvested for analysis.

### siRNA Transfection of Cardiomyocytes

Two siRNA sequences targeting PD-L1 (siPD-L1) and a negative control siRNA (siNC) were synthesized. The sequences for siPD-L1 were 5’-GGAGATTAGATCCTGAGGA-3’ and 5’-CCATACAACAAAATCAACC-3’.

AC16 cells were seeded into six-well plates and transiently transfected with siRNA using the riboFECT™ CP Reagent Kit (Ribobio, Shanghai, China). Forty-eight hours post-transfection, cells were harvested, and PD-L1 mRNA and protein expression levels were assessed to confirm knockdown efficiency.

### Preparation of Human Peripheral Blood Effector Cells

Peripheral blood samples were obtained from healthy adult volunteers under a protocol approved by the Biomedical Ethics Committee of Hebei University of Engineering (Approval No. BER-YXY-2024023). Written informed consent was obtained from all participants before blood collection. PBMCs were isolated through density-gradient centrifugation. CD8⁺ T cells were enriched using magnetic bead sorting (Miltenyi, Order No. 130-045-201), yielding isolated CD8⁺ T cells and CD8⁺ T cell-depleted PBMCs. Enriched CD8⁺ T cells were either activated using anti-CD3/CD28 beads (Miltenyi, Order No. 130-128-758) or maintained without activation for 24 h in RPMI-1640 medium at 37°C with 5% CO₂. PBMCs/CD8⁺ T cells from three independent donors were used.

### Co-culture of Effector Cells and Human Cardiomyocytes

AC16 cells, either irradiated or non-irradiated, were seeded in 96-well plates at 4 × 10³ cells/well. PBMCs, CD8⁺ T-cell-depleted PBMCs, activated CD8⁺ T cells, or non-activated CD8⁺ T cells were subsequently added at the indicated effector-to-target (E:T) ratios (0:1, 5:1, or 10:1). Camrelizumab was added at final concentrations of 0, 0.5, or 5 μg/mL. In separate experiments, irradiated or non-irradiated AC16 cells transfected with siNC or siPD-L1 were co-cultured with activated CD8⁺ T cells at the indicated E:T ratios. Co-cultures were maintained for 24–72 h as specified for each experiment.

### Western Blot

AC16 cells were harvested at the indicated time points after irradiation and/or siRNA transfection. Total protein was extracted using lysis buffer, and protein concentrations were determined using a BCA assay. Equal amounts of protein were separated by SDS-PAGE and transferred onto PVDF membranes. After blocking with 5% BSA for 1 h, membranes were incubated overnight at 4°C with primary antibodies against PD-L1 (Proteintech, Cat. No. 66248-1-Ig, 1:5000), PD-L2 (Proteintech, Cat. No. 18251-1-AP, 1:1000), and GAPDH (Proteintech, Cat. No. 81640-5-RR, 1:10000).

After washing, membranes were incubated with the appropriate HRP-conjugated secondary antibodies for 2 h at room temperature. Protein bands were visualized using enhanced chemiluminescence (ECL) substrate and quantified using ImageJ. PD-L1 and PD-L2 expression levels were normalized to GAPDH.

### qRT-PCR Analysis

Total RNA was extracted from AC16 cells using TRIzol reagent and reverse-transcribed into cDNA using MonScript™ RTIII Super Mix with dsDNase (Two-Step) (Cat. No. MR05201, Monad Biotech, Suzhou, China) according to the manufacturer’s instructions. Quantitative real-time PCR was performed using an ABI 5700 system with SYBR Green reagent. GAPDH was used as the internal reference. Relative mRNA expression was calculated using the 2−ΔΔCt method. Primer sequences are provided in Supplementary Table 1.

### Immunofluorescence

AC16 cells were seeded in confocal culture dishes at a density of 5 × 10³ cells per dish and cultured for 24 h before irradiation. Following 10 Gy irradiation or sham treatment, cells were processed for immunofluorescence at 24, 48, and 72 h. After exposure to 10 Gy irradiation or sham treatment, cells were washed with PBS, fixed with 4% paraformaldehyde (PFA) for 15 min at room temperature. After washing three times with PBS, cells were permeabilized with 0.3% Triton X-100 in PBS for 5 min, followed by blocking with 5% BSA in PBS for 1 h at room temperature. Cells were incubated overnight at 4°C with primary antibodies against PD-L1 (Proteintech, Cat. No. 66248-1-Ig, dilution 1:200) and PD-L2 (Proteintech, Cat. No. 27406-1-AP, dilution 1:300). After three washes with PBS, cells were incubated with the appropriate fluorophore-conjugated secondary antibodies (Proteintech, Cat. Nos. SA00013-1 and SA00013-4, dilution 1:100 and 1:250) for 1 h at room temperature in the dark. Nuclei were counterstained with DAPI. Fluorescence images were acquired using a Nikon A1R confocal microscope with appropriate fluorescence channels for the respective fluorophores.

### Cell Viability Assay

AC16 cells subjected to irradiation and siRNA-mediated PD-L1 knockdown were seeded at 4 × 10³ cells/well in 96-well plates, incubated for 24–72 h, and treated with CCK-8 reagent. Before CCK-8 measurement, non-adherent immune cells were removed by gentle washing with PBS, fresh culture medium was added, and AC16 cell viability was subsequently assessed using the CCK-8 assay. OD values at 450 nm were measured, and the assay was similarly conducted for co-cultured cell groups. Each experiment was independently repeated three times, and cell viability was calculated using the formula:

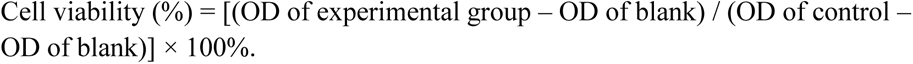

### ELISA

AC16 cells, both irradiated and siRNA-transfected for PD-L1 knockdown, were cultured in six-well plates at 2.5 × 10⁵ cells/well for 24–72 h. Supernatants were collected, and concentrations of IL-6, IL-10, CCL5, CXCL10, and MCP-1 were quantified using ELISA kits (Shanghai Enzyme-linked Biotechnology Co., Ltd.). Standard curves were employed for quantification, and all experiments were independently repeated three times.

### Flow Cytometry Analysis (FACS)

For surface PD-L1 detection, AC16 cells were harvested at the indicated time points after irradiation and/or siRNA transfection, washed with PBS, and incubated with fluorophore-conjugated anti-human PD-L1 antibody (BioLegend, USA, Cat. No. 329705) for 30 min at 4°C in the dark. For PD-1 analysis, CD8⁺ T cells were stained with anti-human PD-1 antibody (BioLegend, USA, Cat. No. 367418). Samples were analyzed using FlowJo v10.6.2.

### Apoptosis Analysis

Apoptosis was assessed using an Annexin V-FITC/PI apoptosis detection kit (Beyotime, Cat. No. C1062S, Shanghai, China) according to the manufacturer’s instructions. AC16 cells were harvested at the indicated time points after irradiation and/or siRNA transfection, washed with PBS, resuspended in binding buffer, and stained with Annexin V and PI for 10 min in the dark. Samples were analyzed using a CytoFLEX LX flow cytometer (Beckman Coulter, USA). Early and late apoptotic cells were defined according to Annexin V/PI staining, and the total apoptotic fraction was calculated as the sum of early-apoptotic and late-apoptotic cell proportions.

### Hematoxylin and Eosin (HE) Staining

Excised heart tissues were fixed, paraffin-embedded, sectioned, and stained with HE to evaluate cellular morphology, with nuclei and cytoplasm visualized in blue and red, respectively.

### Wheat Germ Agglutinin (WGA) Staining

Cardiomyocyte morphology and size were assessed in mouse heart sections using WGA immunofluorescence staining. Cardiomyocytes appeared green, and nuclei were stained blue. Sections were blocked with goat serum and incubated sequentially with primary antibodies (1:100), secondary antibodies (1:100), and WGA stain (1:100) in the dark, followed by DAPI mounting and imaging.

### Immunofluorescence Staining

Frozen mouse heart sections (20–30 μm) were rinsed with PBS, permeabilized, and blocked with PBS containing Triton X-100 and goat serum for one hour. Sections were incubated overnight at 4°C with primary antibodies against PD-L1, CD8, and CD4, washed, and then treated with fluorescent secondary antibodies for 2 hours in the dark. Slides were mounted with antifade medium and observed under a confocal microscope with the following excitation/emission settings: DAPI (330–380 nm/420 nm), Alexa Fluor 488 (465–495 nm/515–555 nm), Cy3 (510–560 nm/590 nm), and Cy5 (608–648 nm/672–712 nm).

### Statistical Analysis

Data were expressed as mean and standard deviation (SD). Comparisons between two groups were conducted using Student’s *t*-test, while one-way analysis of variance (ANOVA) with Bonferroni’s post-hoc test was applied for comparisons involving three or more groups. For experiments involving two independent factors, two-way ANOVA followed by Tukey’s multiple comparisons test was used. Statistical significance was defined as *p* < 0.05. Data analysis was performed using GraphPad Prism 9, ImageJ-win64, FlowJo v10.6.2, and EZ-C1 software. Figures were prepared using Adobe Illustrator.

## Results

### Irradiation induces time-dependent PD-L1 upregulation in AC16 cells

To elucidate the regulation of PD-L1 expression in cardiomyocytes after irradiation, AC16 cells were exposed to 6 Gy and 10 Gy irradiation, then cultured for 24 to 72 hours post-irradiation. Flow cytometry showed that membrane PD-L1 expression was significantly increased after 10 Gy irradiation compared with controls (29.06% ± 4.437 vs. 6.45% ± 1.544), with a progressive increase over time (Fig. 1A, B). No significant changes were observed in the 6 Gy group (Fig. 1A, B). Western blot analysis showed increased PD-L1 protein expression with consistently low PD-L2 expression pre- and post-irradiation. qRT-PCR analysis confirmed significant up-regulation of PD-L1 mRNA at 48 hours following 10 Gy irradiation (Fig. 1E). Immunofluorescence imaging indicated a translocation in PD-L1 localization from even distribution in unirradiated cells to cytoplasmic accumulation at 24 and 48 hours post-irradiation. PD-L1 fluorescence was predominantly observed in membrane- and cytoplasm-associated regions at 24–48 h after irradiation, whereas prominent nuclear-associated fluorescence was observed at 72 h. (Fig. 1F).

**Fig. 1.**
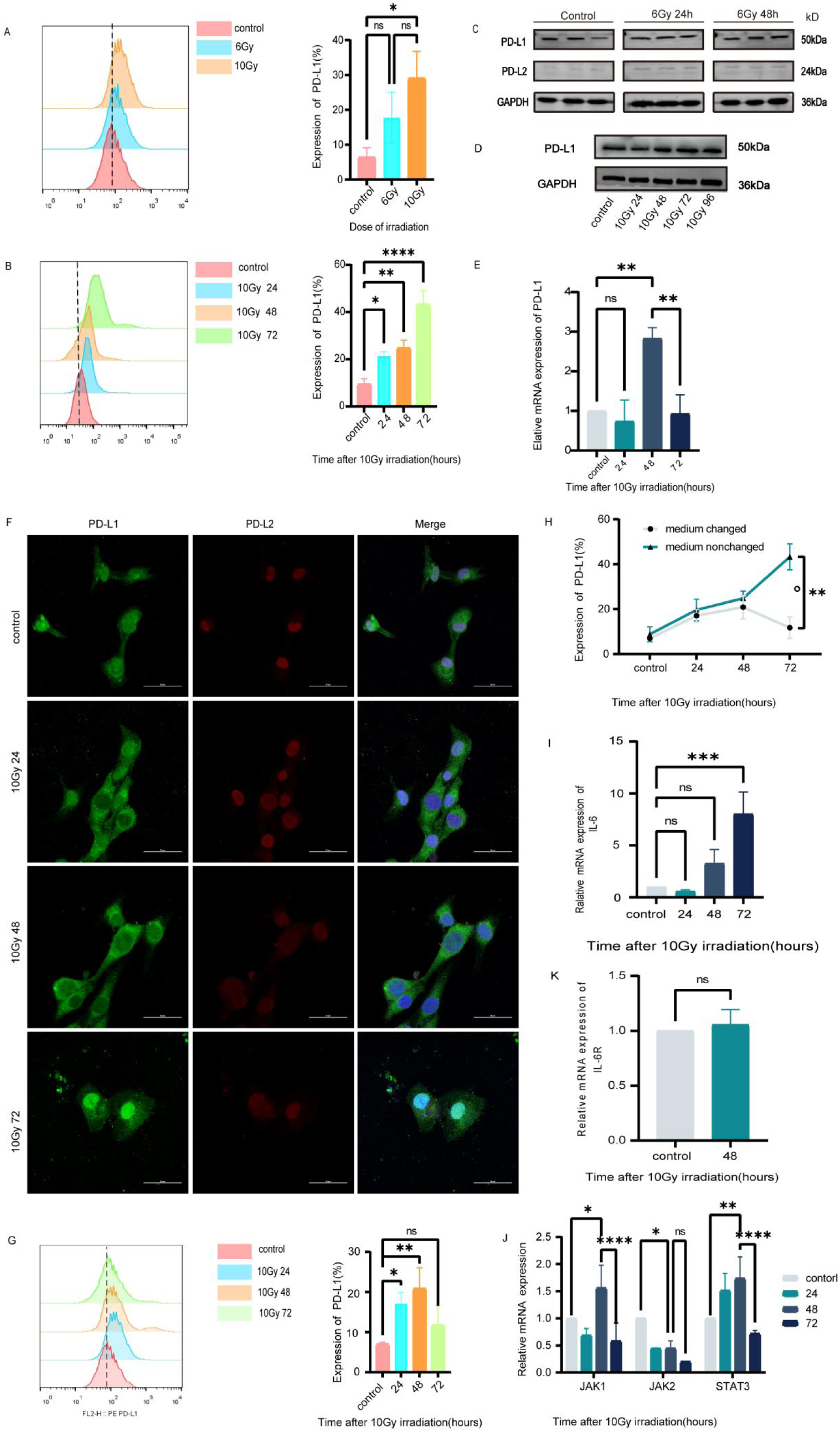
Irradiation induces time-dependent PD-L1 upregulation in AC16 cells. (A) Surf ace PD-L1 expression in AC16 cells was assessed by flow cytometry at 72 h after different irradiation doses. (B) PD-L1 membrane expression was time-dependent after 10 Gy irradiation. (C) The PD-L1 and PD-L2 protein expression was assessed by western blotting at 24 and 48 h after 6 Gy irradiation. (D) PD-L1 protein expression was time-dependent after 10 Gy irradiation. (E) Relative PD-L1 mRNA expression was assessed by qRT-PCR at 24, 48, and 72 h after 10 Gy irradiation. (F) Immunofluorescence analysis of PD-L1 and PD-L2 in AC16 cells at the indicated time points after 10 Gy irradiation. Primary antibodies against PD-L1 were detected using the appropriate fluorophore-conjugated secondary antibodies (Proteintech, Cat. Nos. SA00013-1 and SA00013-4). PD-L1 fluorescence increase d in membrane- and cytoplasm-associated regions at 24–48 h and became prominent in the nuclear region at 72 h. In A, B, and E, the data are presented as the mean ± SD of three independent experiments. (G) PD-L1 membrane expression decreased at 72 h when rep lacing the culture medium of cardiomyocytes at 48 hours after 10 Gy irradiation. (H) PD-L1 membrane expression in the culture replacement group significantly decreased at 72 h after irradiation. (I) The relative mRNA expression level of IL-6 after 10 Gy irradiation. (J) JAK1 and STAT3 mRNA expression increased after irradiation, whereas JAK2 mRNA expression did not change significantly. (K) IL6R mRNA expression was assessed by qRT -PCR before and after irradiation. The data shown are the mean ± SD of three independent experiments. *p < 0.05, **p < 0.01, ***p < 0.001, ****p < 0.0001.

Interestingly, we found that the persistence of up-regulation of membrane PD-L1 expression in the 10 Gy group was influenced by medium conditions. Continuous culture without medium replacement for 24 to 72 hours showed a sustained increase in PD-L1 expression (Fig. 1H), whereas medium replacement at 48 hours resulted in reduced PD-L1 expression at 72 hours (Fig. 1G). Further qRT-PCR analysis indicated significant IL-6 mRNA upregulation post-irradiation (1 vs. 8.029 ± 1.020, p = 0.0006) (Fig. 1I), accompanied by increased JAK1 and STAT3 mRNA. However, JAK2 did not show a sustained increase (Fig. 1J), and IL6R mRNA remained unchanged (IL6R Ct values: control vs. irradiation, 23.60 ± 0.1527 vs. 23.68 ± 0.1232; p > 0.05) (Fig. 1K).

### Irradiation induces AC16 cell injury and alters inflammatory cytokine and chemokine release

AC16 cells were exposed to 10 Gy irradiation and subsequently cultured for 24–72 h to determine apoptosis, cytokine/chemokine concentrations, and cell viability through flow cytometry, ELISA, and CCK-8 assays. Apoptosis rates were significantly higher in cardiomyocytes exposed to 10 Gy irradiation at 48 and 72 hours post-irradiation (17.47% ± 1.684 and 22.72%±1.130, respectively; *p* < 0.001), compared with the control group (8.353%±1.689) (Fig. 2A). A similar but attenuated trend was observed in the 6 Gy group (Fig. 2A). On the basis of these findings, 10 Gy was selected for subsequent experiments. Cell viability, as assessed by CCK-8 assays, was markedly reduced at 48 and 72 hours post-irradiation in the irradiated (IR+) group relative to the non-irradiated (IR-) group (Fig. 2B). ELISA demonstrated time-dependent increases in IL-6, MCP-1, CCL5, and CXCL10, accompanied by a decrease in IL-10, in the supernatants of irradiated AC16 cells compared with non-irradiated controls (Fig. 2C and Supplementary Table 2).

**Fig. 2.**
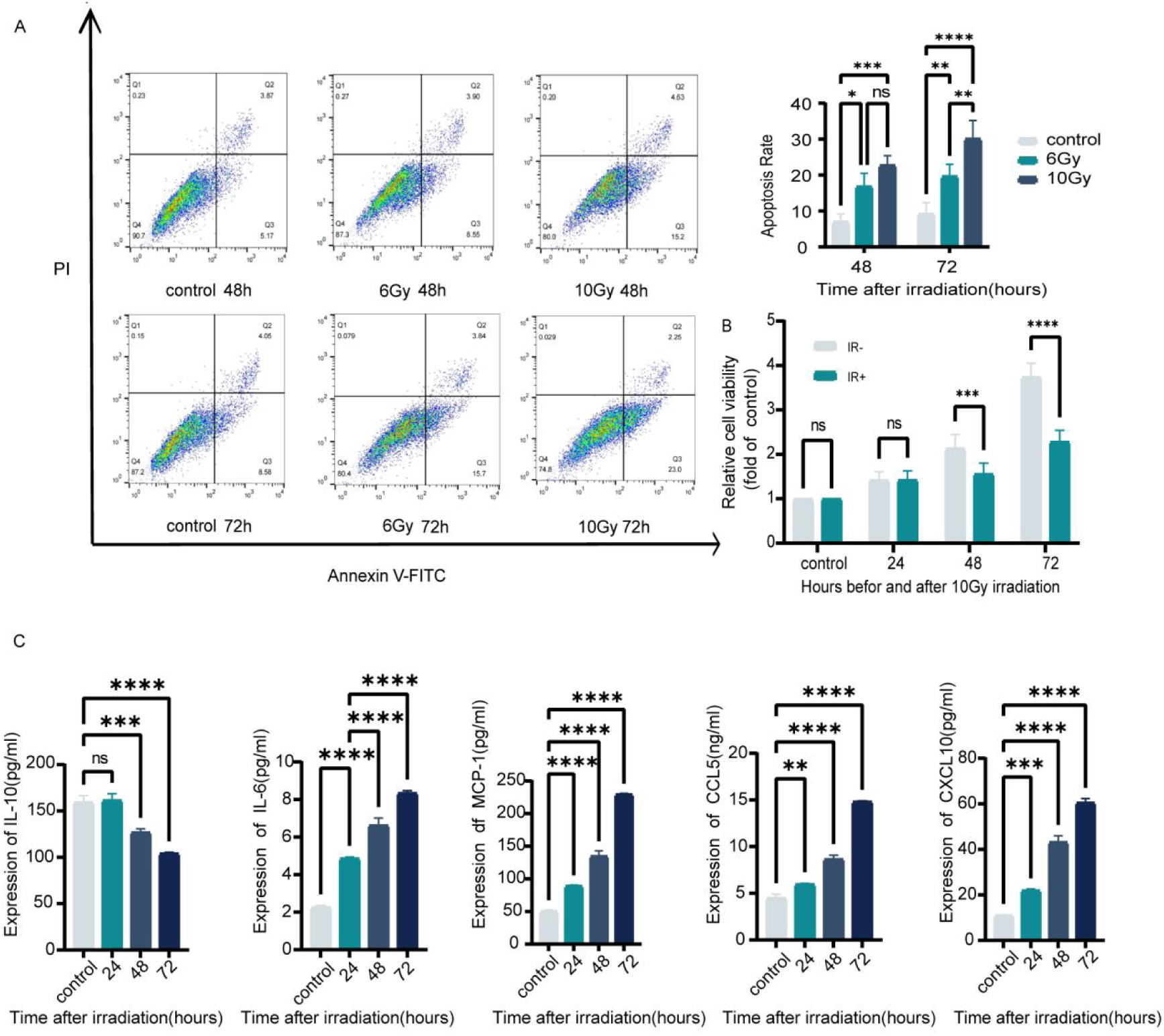
Irradiation induces AC16 cell injury and alters inflammatory cytokine and chemokine release. (A) Apoptosis rates were assessed by flow cytometry at 48h and 72 h after 6 or 10 Gy irradiation. (B) Comparison of cardiomyocyte viability before and after 10 Gy irradiation at 24h, 48h, and 72 h. (C) ELISA was used to quantify IL-10, IL-6, MCP-1, CCL5, and CXCL10 concentrations in culture supernatants before and after 10 Gy irradiation. Data are presented as mean ± SD of three independent experiments. *p < 0.05, **p < 0.01, ***p < 0.001, ****p < 0.0001.

### PD-L1 knockdown enhances inflammatory cytokine and chemokine responses in AC16 cells

To explore the effect of PD-L1 in cardiomyocyte homeostasis and function, PD-L1 expression was silenced using siRNA (Supplementary Fig. 1). Then apoptosis, cytokine/chemokine concentration, and cell viability were examined by the same methodologies. PD-L1 mRNA expression was significantly down-regulated at 48 hours after irradiation (qRT-PCR, Fig. 3A) and PD-L1 protein expression in AC16 cells at 72 hours (western blotting and flow cytometry, Fig. 3B, C) compared with non-knockdown controls. The knockdown of PD-L1 did not significantly alter apoptosis rates in either irradiated (IR+) or non-irradiated (IR-) groups (*p*>0.05, Fig. 3D). However, it markedly affected cell viability and cytokine/chemokine concentration. ELISA results showed significantly higher concentrations of IL-6, CCL5, and CXCL10 in the siPD-L1 group compared with siNC controls, with the effect more pronounced in the non-irradiated groups (Fig. 3E, Supplementary Tables 3 and 4). Furthermore, CCK-8 assays revealed a significant reduction in cell viability in the siPD-L1 group compared with siNC under non-irradiated conditions, while no significant difference was observed in irradiated cells (Fig. 3F).

**Fig. 3.**
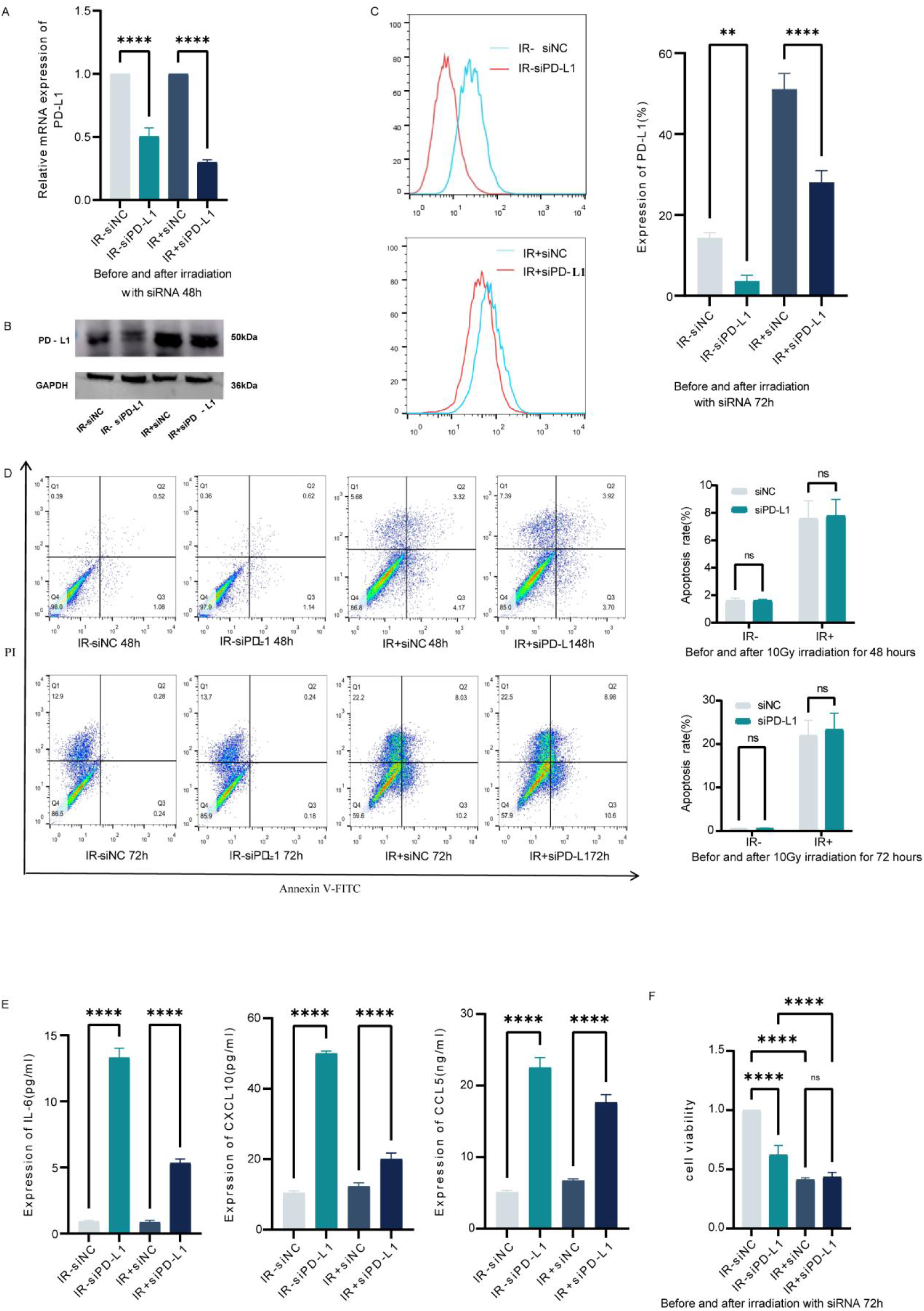
PD-L1 knockdown enhances inflammatory cytokine and chemokine responses in AC16 cells. (A) qRT-PCR was used to verify the knockdown efficiency of PD-L1 at the mRNA level. (B) Western blotting confirmed the knockdown efficiency of PD-L1 at the protein level. (C) Flow cytometry was used to assess the knockdown efficiency of surface PD-L1 expression. (D) Flow cytometry compared apoptosis in irradiated and non-irradiated cardiomyocytes before and after PD-L1 knockdown. (E) ELISA analyzed differences in IL-6, CCL5, and CXCL10 concentrations in irradiated and non-irradiated cardiomyocytes before and after PD-L1 knockdown. (F) CCK-8 assays compared the cell viability of irradiated and non-irradiated cardiomyocytes before and after PD-L1 knockdown. Data are presented as mean ± SD of three independent experiments. *p < 0.05, **p < 0.01, ***p < 0.001, ****p < 0.0001.

### The protective role of the PD-1/PD-L1 pathway in iRT-induced cardiomyocyte injury

An in vitro model was established to examine the mechanisms underlying iRT-induced cardiomyocyte injury (Fig. 4A). Human PBMCs were co-cultured with AC16 cells both pre- and post-irradiation at an effector-to-target (E:T) ratio of 10:1 with and without ICI (5 μg/mL), and incubated for 24 h. AC16 cell viability was lowest in the IR+ICI group (p < 0.01) (Fig. 4B). These viability differences were not observed when CD8^+^ T-cell-depleted PBMCs or non-activated CD8^+^ T cells were used, whereas co-culture with activated CD8⁺ T cells significantly reduced AC16 cell viability (Fig. 4C, D).

**Fig. 4.**
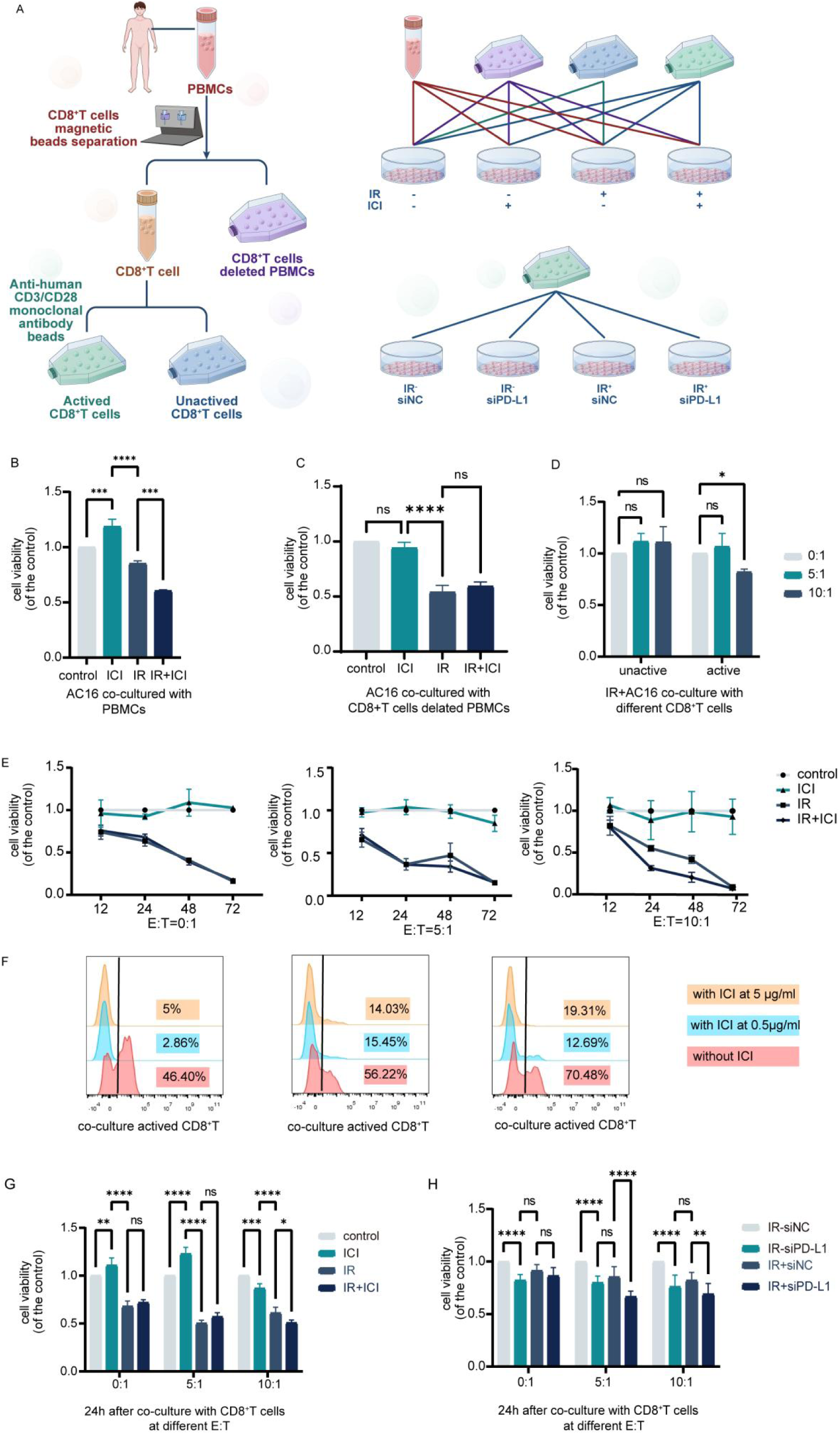
PD-1/PD-L1 signaling modulates activated CD8⁺ T-cell-associated injury in AC16 co-culture. (A) Schematic of the preparation of human peripheral blood effector cells and co-culture with human AC16 cells. (B) PBMCs were co-cultured with irradiated and non-irradiated cardiomyocytes at an E:T ratio of 10:1, with the addition of ICI (5 μg/mL), and CCK-8 was used to assess cell viability in four groups of cardiomyocytes. (C) After removing CD8⁺ T cells, CD8^+^ T-cell-depleted PBMCs were co-cultured with irradiated and non-irradiated cardiomyocytes at an E:T ratio of 10:1, with ICI concentrations of 5 μg/mL, and CCK-8 was used to assess cardiomyocyte survival. (D) Activated and non-activated CD8⁺ T cells were co-cultured with irradiated cardiomyocytes at an E:T ratio of 5:1 or 10:1, and CCK-8 was used to assess cell survival. (E) Activated CD8⁺ T cells at different effector-to-target ratios (E:T ratios of 0:1, 5:1, and 10:1) were co-cultured with irradiated and non-irradiated cardiomyocytes, with or without ICI (5 μg/mL), for 12-72 h, and CCK-8 was used to assess AC16 cell viability over time. (F) Flow cytometry was used to test detectable surface PD-1 staining on CD8⁺ T cells after treatment with different concentrations (0, 0.5, and 5 μg/mL) of anti-PD-1 antibody. (G) Activated CD8⁺ T cells were co-cultured with irradiated and non-irradiated cardiomyocytes at an E:T ratio of 10:1 for 24 hours in four groups: control group (control), PD-1 monoclonal antibody group (ICI group), irradiation group (IR group), and irradiation combined with PD-1 monoclonal antibody group (IR+ICI). CCK-8 was used to assess AC16 cell viability. (H) Activated CD8⁺ T cells were co-cultured with PD-L1 knockdown and non-knockdown cardiomyocytes at an E:T ratio of 10:1 for 24 hours in four groups: control co-culture group (IR-siNC), PD-L1 knockdown co-culture group (IR-siPD-L1), irradiation co-culture group (IR+siNC), and irradiation combined with PD-L1 knockdown co-culture group (IR+siPD-L1), and CCK-8 was used to assess AC16 cell viability in each group. Data are presented as the mean ± SD of three independent experiments. *p < 0.05, **p < 0.01, ***p < 0.001, ****p < 0.0001. The camrelizumab concentration used in panel G was 0.5 μg/mL.

Subsequently, activated CD8^+^ T cells were co-cultured with AC16 cells both pre- and post-irradiation under varying E:T ratios (0:1, 5:1, 10:1) and ICI concentrations (0, 0.5, 5 μg/mL), and incubated for 12–72 h. At an E:T ratio of 10:1 and an ICI concentration of 5 μg/mL, after 24 and 48 h of incubation, AC16 cell viability was lowest in the IR+ICI group (p < 0.01). However, this trend was not observed at an E:T ratio of 5:1or at incubation times of 12 h and 72 h (Fig. 4E) (Supplementary Fig. 2 and 3).

To investigate PD-1/PD-L1 pathway involvement, PD-1 on CD8⁺ T cells was blocked using ICI (Fig. 4F), and PD-L1 in AC16 cells was knocked down via siRNA. The proportions of activated CD8^+^ T cells with detectable surface PD-1 staining were 57.7% ± 12.11 without ICI, 10.33% ± 3.82 with 0.5 μg/mL ICI, and 12.78% ± 4.18 with 5 μg/mL ICI. No significant difference was observed between the 0.5 and 5 μg/mL groups (Fig. 4F and Supplementary Fig. 5). The co-culture results indicate that, at an E:T ratio of 10:1 and an ICI concentration of 0.5 μg/mL, disruption of PD-1/PD-L1 signaling was associated with a further reduction in AC16 cell viability (Fig. 4G, H).

### Pathological observations of myocardial injury in vivo

An in vivo model of iRT-induced myocardial injury was successfully established (Fig. 5A). HE staining showed orderly myocardial cell alignment, clear structures, and centrally located nuclei in control, with no discernible abnormalities. In the ICI group, mild structural disorganization was observed, including slight edema, vacuolization in some cells, and minimal inflammatory cell infiltration. The IR group exhibited more pronounced disorganization, including significant cellular and stromal edema, partial myolysis, nuclear condensation and displacement, and focal inflammatory infiltration. In the IR+ICI group, focal regions demonstrated pronounced edema, myocardial fiber rupture with myolysis, while structural damage did not substantially exceed that of the IR group. CD8^+^ T-cell infiltration appeared somewhat greater in the IR+ICI group than in the other groups but remained limited and focal. No widespread myocardial necrosis was observed, and no mice died during the 28-day observation period (Fig. 5D). WGA staining revealed an increased myocardial cell cross-sectional area in the ICI, IR, and IR+ICI groups compared with controls (Fig. 5C). Immunofluorescence analysis confirmed lymphocyte infiltration in the ICI, IR, and IR+ICI groups, CD8⁺ T cell infiltration was notably greater than CD4⁺ T cell infiltration (Fig. 5B). PD-L1 immunofluorescence staining demonstrated low expression in the control group, with markedly increased expression in the ICI, IR, and IR+ICI groups (Fig. 5E). Dual staining for PD-L1 and WGA further revealed elevated PD-L1 expression on both the cell membrane and within the cytoplasm of myocardial cells in treated groups relative to controls (Fig. 5F).

**Fig. 5.**
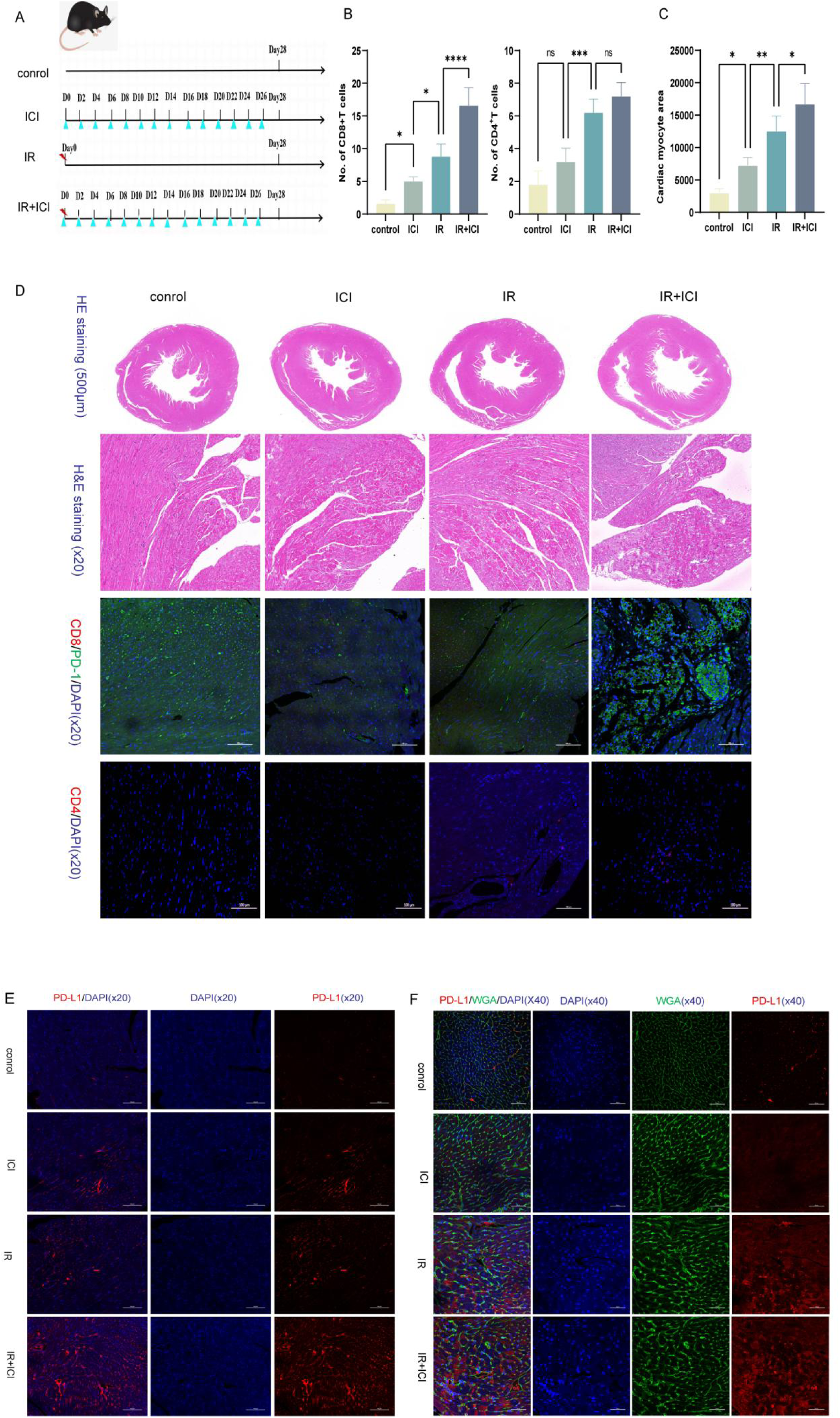
Histopathological and immunofluorescence assessment of myocardial injury, T-cell infiltration, and PD-L1 expression in vivo. (A) Schematic of irradiation (IR) and/or PD-1 inhibitor (ICI) treatment in C57BL/6 mice. Mice received localized 20 Gy cardiac irradiation and/or anti-PD-1 antibody treatment as indicated. (B) CD8⁺ and CD4⁺ T-cell infiltration in myocardial tissue was assessed by immunofluorescence. (C) Quantitative analysis of cardiomyocyte cross-sectional area using WGA staining. (D) Hematoxylin and eosin (HE) staining of myocardial tissue at day 28 after irradiation and/or PD-1 blockade. (E) Myocardial PD-L1 expression assessed by immunofluorescence. (F) Dual staining of PD-L1 and WGA showing PD-L1 distribution in myocardial tissue. Control, untreated control; ICI, PD-1 inhibitor; IR, irradiation; IR+ICI, irradiation plus PD-1 inhibitor.

**Fig. 6.**
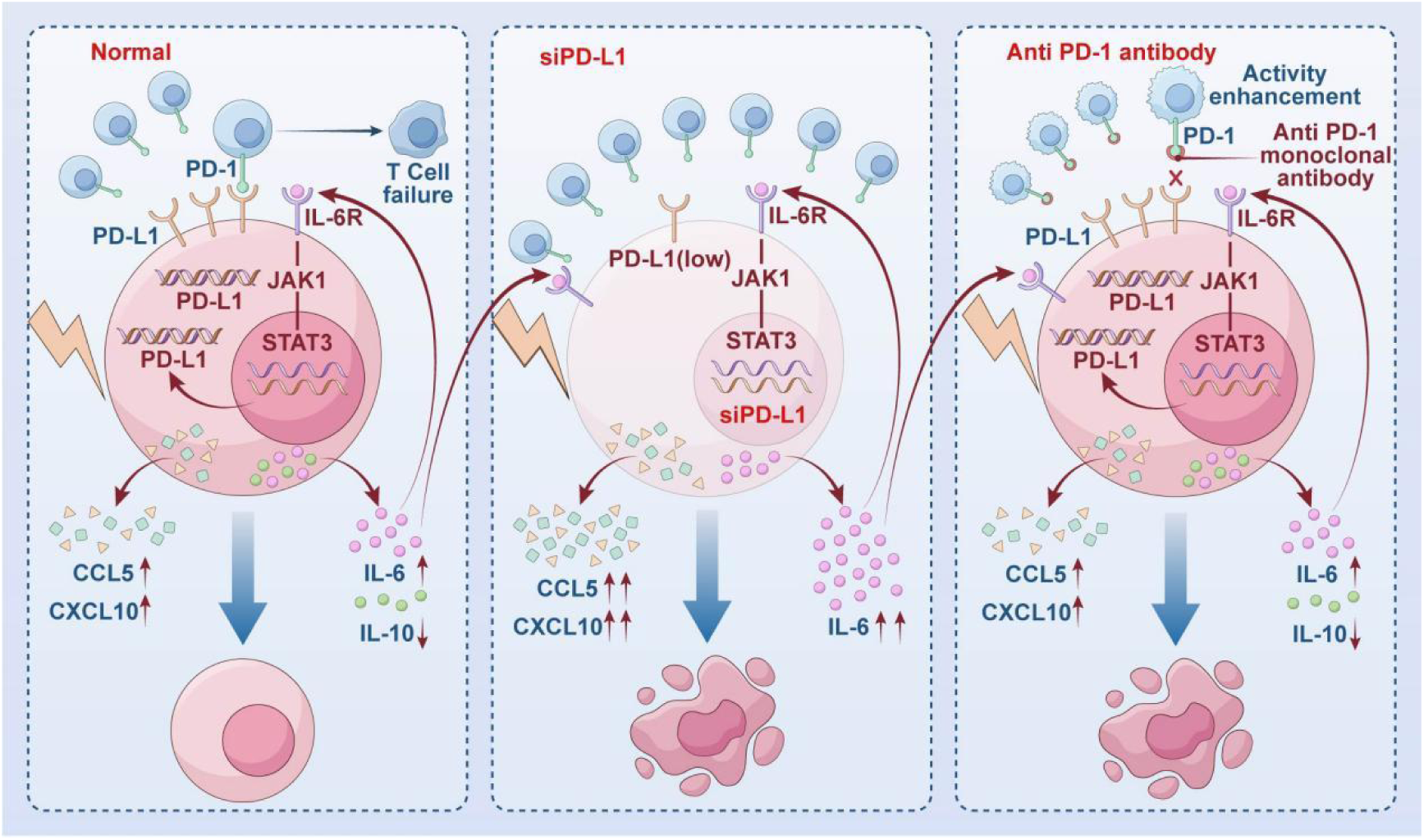
Proposed working model of radiation-induced cardiomyocyte PD-L1 regulation and its potential immunoregulatory role during iRT-associated injury. Irradiation induces PD-L1 expression in AC16 cells while simultaneously increasing the release of inflammatory cytokines and chemokines. PD-L1 knockdown further increases IL-6, CCL5, and CXC L10 release, suggesting that PD-L1 contributes to inflammatory homeostasis. In the presence of abundant activated CD8⁺ T cells, disruption of PD-1/PD-L1 signaling by PD-1 blockade or PD-L1 knockdown is associated with reduced AC16 cell viability. IL-6/JAK1/STAT3 signaling is proposed as a potential upstream regulator of radiation-induced PD-L1 expres sion. This proposed pathway is hypothesis-generating and requires further experimental vali dation.

## DISCUSSION

The combination of radiotherapy and anti-PD-1 antibody has demonstrated significant survival benefits in patients with thoracic tumors. However, whether these combined therapies increase the risk of cardiac toxicity remains uncertain ^[24,25]^. The phase III ADRIATIC trial showed that durvalumab after concurrent chemoradiotherapy significantly improved overall survival and progression-free survival compared with placebo in patients with limited-stage small-cell lung cancer, with similar rates of grade 3–4 adverse events (24.4% vs. 24.2%) ^[24]^. A pooled analysis by Anscher et al. found no meaningful increase in serious adverse events among patients who received an ICI within 90 days after radiotherapy ^[9]^. Conversely, Du et al. indicated that combining irradiation with anti-PD-1 antibody treatment in mice resulted in an acute mortality rate of 30% within 2 weeks, significantly higher than the 0% mortality observed in mice receiving cardiac irradiation plus control IgG ^[13]^. These discordant findings between clinical and preclinical studies highlight the need for further investigation into the mechanisms underlying myocardial injury induced by iRT.

Several studies have separately explored the mechanisms of radiation-induced heart disease and ICI-associated cardiotoxicity ^[26–28]^, but mechanistic studies of cardiac injury induced by combined radiotherapy and immune checkpoint inhibition remain limited. In the present study, we established an in vitro model to investigate the response of human cardiomyocytes to irradiation in the context of PD-1/PD-L1 signaling. Irradiation induced PD-L1 upregulation together with increased inflammatory cytokine and chemokine release. These observations suggest that cardiomyocytes may activate an adaptive immunoregulatory response during radiation-associated inflammatory stress. In vivo, myocardial PD-L1 expression was also increased, whereas CD8⁺ T-cell infiltration remained limited and focal.

Our findings suggest that radiation-induced PD-L1 upregulation in cardiomyocytes may represent an adaptive immunoregulatory response. After irradiation, cardiomyocyte viability decreased, accompanied by increased levels of inflammatory cytokines (e.g., IL-6 and MCP-1) and chemokines (e.g., CCL5 and CXCL10). Interestingly, PD-L1 knockdown produced distinct effects on inflammatory regulation and cardiomyocyte viability. Silencing PD-L1 markedly increased the release of IL-6, CCL5, and CXCL10 under both irradiated and non-irradiated conditions, supporting a role for PD-L1 in restraining inflammatory and chemokine responses. In contrast, PD-L1 knockdown reduced cell viability under non-irradiated conditions but did not further decrease viability in irradiated cardiomyocytes, and apoptosis was not significantly altered. These findings suggest that the protective function of cardiomyocyte PD-L1 may not primarily reflect a direct anti-apoptotic effect against radiation injury. Instead, PD-L1 may function as an adaptive immunoregulatory mechanism that helps maintain cellular and inflammatory homeostasis. The functional importance of this adaptive PD-L1 response became more apparent in the presence of activated CD8⁺ T cells. Under high effector-to-target conditions, disruption of PD-1/PD-L1 signaling by PD-1 blockade or PD-L1 knockdown further reduced cardiomyocyte viability, suggesting that cardiomyocyte PD-L1 may exert a prominent protective effect in an immune-cell-dependent context.

Additionally, our experiments demonstrated that when disrupting the PD-1/PD-L1 pathway via anti-PD-1 antibodies or siRNA-mediated PD-L1 knockdown, the AC16 cardiomyocyte viability in iRT group was significantly decreased when compared with irradiation group or anti-PD-1 antibody group alone after co-culturing these cells with abundant activated CD8⁺ T cells. However, we observed that co-culturing with a limited density of activated CD8⁺ T cells or non-activated CD8⁺ T cells, cardiomyocyte viability was not significantly reduced. In vivo, myocardial PD-L1 expression was increased after iRT, whereas CD8⁺ T-cell infiltration remained limited and focal, and no deaths were observed during the 28-day observation period. This pattern is directionally consistent with some clinical observations showing no prominent increase in fatal cardiotoxicity with combined radiotherapy and immune checkpoint inhibition. Together with our in vitro findings, these observations raise the possibility that adaptive cardiac immune-regulatory responses may partly contribute to the discrepancy between some preclinical models and clinical observations. However, the absence of mortality during the 28-day observation period does not exclude subclinical or delayed cardiac injury.

Although PD-L1 expression in cardiomyocytes has increasingly been implicated in cardiac immune homeostasis, the mechanisms regulating radiation-induced PD-L1 upregulation in cardiomyocytes remain poorly understood. Our study demonstrated that irradiation induces significant PD-L1 upregulation in cardiomyocytes, with distinct spatiotemporal changes in its subcellular distribution. PD-L1 fluorescence was predominantly observed in membrane- and cytoplasm-associated regions at earlier time points, whereas prominent nuclear-associated fluorescence was observed at 72 h. This observation parallels the findings in breast cancer cells, in which radiation-induced nuclear translocation of PD-L1 was observed, and the phenomenon is associated with pyroptosis ^[29]^. Whether similar mechanisms exist in human cardiomyocytes remains to be determined. Current considerations of the mechanism of PD-1 and PD-L1 inhibitors focus their effects on T cells with membrane-expressed PD-1, while with limited understanding of the effects of PD-L1 in the cytoplasm and nucleus; its short cytoplasmic tail lacks a classical signaling motif, although noncanonical intracellular or reverse-signaling functions have been proposed ^[30]^. Accordingly, the spatiotemporal changes in PD-L1-associated fluorescence observed in AC16 cells warrant further investigation using complementary subcellular localization approaches.

In our study, PD-L1 upregulation in cardiomyocytes after irradiation was attenuated by replacing the culture medium, suggesting that sustained PD-L1 elevation may be influenced by soluble mediators released from irradiated cells. Previous studies have shown that IL-6, IFN-γ, and TNF-α are mediators of PD-L1 expression ^[30,31]^. Our findings showed that irradiation increased IL-6 release together with JAK1 and STAT3 mRNA expression, whereas JAK2 and IL6R mRNA levels were not significantly altered. Sasi et al. showed that IL-6 can participate in an autocrine loop that sustains PD-L1 expression through JAK/STAT signaling in normal and malignant lymphocytes ^[32]^. More directly, Zhou et al. demonstrated radiotherapy-mediated IL-6/STAT3-dependent PD-L1 upregulation in esophageal cancer cells ^[33]^. These studies provide biological plausibility for an IL-6/JAK1/STAT3–PD-L1 axis in irradiated cardiomyocytes. Whether an IL-6/JAK1/STAT3-PD-L1 axis operates in irradiated cardiomyocytes remains to be established ^[34]^, but our current data do not establish pathway activation or causality; this hypothesis requires direct experimental validation.

Identifying reliable biomarkers is crucial for the early detection of cardiomyocyte injury. Previous research has highlighted that CCL5 and CXCL10 are inflammatory chemokines implicated in T-cell recruitment and inflammatory cardiac injury. Spatial transcriptomic analysis of viral myocarditis identified CXCL9/CXCL10-rich inflamed regions associated with cytotoxic T-cell recruitment ^[35]^. A study using multiomics single-cell technology to identify immune cell subsets associated with ICI-induced myocarditis revealed that the elevated levels of proinflammatory chemokines, including CCL5, in clonally expanded CD8⁺ T cells, were linked to fatal myocarditis. ^[36]^. Sildenafil reduced CXCL10 expression and secretion in stimulated human cardiomyocytes and lowered circulating CXCL10 in a subgroup of patients with diabetic cardiomyopathy, suggesting potential modulation of CXCL10-associated inflammation ^[37]^. Evidence remains limited on the correlation between these two cytokines and RIHD. Consistent with these findings, our study revealed that after irradiation, the release of CCL5 and CXCL10 from cardiomyocytes increased significantly. After PD-L1 knockdown, CCL5 and CXCL10 secretion increased further. These findings identify CCL5 and CXCL10 as candidate inflammatory mediators associated with PD-L1 disruption; their potential value as biomarkers requires clinical validation.

Experimental myocarditis models show that disruption of PD-1/PD-L1 signaling can amplify T-cell-mediated cardiac inflammation. PD-1-deficient MRL mice develop fatal myocarditis with massive CD4⁺ T and CD8⁺ T-cell infiltration ^[19]^, whereas PD-L1 deficiency worsens autoimmune myocarditis in MRL mice ^[17]^. PD-1 signaling has also been shown to limit T-cell-mediated myocardial inflammation and myocyte damage ^[38]^. Previous studies have demonstrated that PD-L1-deficient mice exhibited an increased mortality due to the abundant CD8⁺ T-cells infiltration in myocardium. In the cMy-mOva myocarditis mouse model, adoptive transfer of OVA-specific CD8⁺ T cells at low dose (25 × 10^3^ OT-I CTLs) resulted in reversible myocardial injury, whereas it would cause lethal myocardial injury at high dose (>5 × 10^5^ OT-I CTLs). In contrast, infiltrating CD4⁺ T cells exhibited no significant cardiotoxicity ^[18,39]^. Our in vitro experiments showed a similar context dependence: AC16 cell viability decreased more with a higher number of activated CD8⁺ T cells (4 × 10⁴) than with a lower number (2 × 10⁴), particularly under iRT conditions, whereas non-activated CD8⁺ T cells or CD8⁺ T-cell-depleted PBMCs produced less effect. These findings indicate that both CD8⁺ T-cell abundance and activation status are important determinants of cardiomyocyte injury. Our findings extend previous evidence implicating CD8⁺ T cells in radiation- and immune checkpoint blockade-associated cardiac injury by showing, in a human cardiomyocyte co-culture system, that the extent of injury varies with CD8⁺ T-cell abundance and activation status. In our C57BL/6 mice, CD8⁺ T-cell infiltration in the iRT group appeared somewhat greater than in the other groups but remained limited and focal, with no widespread myocardial necrosis or death.

Several limitations should be acknowledged. First, the in vitro experiments were primarily performed using the AC16 cell line and relatively high single irradiation doses, which do not fully reproduce clinically fractionated cardiac irradiation. Second, cardiomyocyte-specific manipulation of PD-L1 was not performed in vivo, limiting causal inference regarding its protective function. Third, the proposed IL-6/JAK1/STAT3 mechanism is based on transcriptional and secreted-factor changes and requires direct validation of pathway activation. Fourth, the nuclear-associated PD-L1 signal requires confirmation using complementary subcellular localization approaches. Finally, the mouse experiments were limited to an early 28-day observation period and therefore do not exclude delayed functional or fibrotic cardiac injury.

In summary, by establishing cardiac injury models induced by iRT both in vivo and in vitro, we demonstrated significant upregulation of PD-L1 expression in human cardiomyocytes after irradiation. This upregulation may contribute to adaptive immunoregulation by restraining inflammatory mediator release. Activated CD8⁺ T cells played a key role in the co-culture model, and the extent of cardiomyocyte injury was associated with their abundance and activation status. Cardiac risk after thoracic radiotherapy is strongly influenced by cardiac dose and may manifest earlier than traditionally recognized, supporting efforts to minimize heart exposure and incorporate cardio-oncology surveillance ^[2]^. Our findings show that the combined-treatment group did not show excess mortality or markedly greater acute histopathological injury than irradiation alone. Although it is consistent with current clinical data, high mortality rate still remains a very important issue for immunotherapy. Noteworthy, improving strategies to prevent radiation-induced cardiac injury remains a critical clinical priority, since recent studies indicated that the impact of cardiac radiation doses on survival might emerge earlier than the long-term effects of radiotherapy on coronary artery disease (CAD) previously observed^[40]^.

## Declarations

### Ethics approval and informed consent

Peripheral blood samples were obtained from healthy adult volunteers under a protocol approved by the Biomedical Ethics Committee of Hebei University of Engineering (Approval No. BER-YXY-2024023). Written informed consent was obtained from all participants before blood collection. All procedures involving human-derived samples were conducted in accordance with institutional guidelines and applicable ethical standards. All experimental procedures involving animals were reviewed and approved by the Animal Welfare and Ethics Committee of the Fourth Hospital of Hebei Medical University (Approval No. 2022035) and were conducted in accordance with institutional guidelines for the care and use of laboratory animals.

### Author contributions

Ye Zhou: Conceptualization, Methodology, Investigation, Formal analysis, Visualization, Writing – original draft.

Yajing Wu: Methodology, Investigation, Validation, Data curation, Animal experiments.

Na Zhang: Investigation, Resources, Data curation, Cell Experiments.

Shuo Wang: Methodology, Visualization, Aniaml experiments.

Xueshuai Ye: Conceptualization, Methodology, Cell co-culture Experiments.

Jingtao Ma: Conceptualization, Supervision, Resources.

Qingxia Li: Writing – review and editing.

Jun Wang: Conceptualization, Supervision, Resources, Funding acquisition, Writing – review and editing.

All authors reviewed and approved the final manuscript.

### Data availability

The data supporting the findings of this study are available from the corresponding author upon reasonable request.

### Prior presentation

Portions of this work were previously presented at the 2024 and 2025 Annual Meetings of the American Society for Radiation Oncology (ASTRO) and were published in abstract form (doi: 10.1016/j.ijrobp.2024.07.908; doi: 10.1016/j.ijrobp.2025.06.3090).

