## supplemental for "Cardiomyocytes upregulated PD-L1 expression to alleviate cardiac injury induced by irradiation combined with anti-PD-1 antibody: an in vitro and in vivo study"

**SupplementalTable 1. Primers used in qRT-PCR**

| Primer | Forward | | | Reverse | |
| --- | --- | --- | --- | --- | --- |
| PD-L1 | 5’-ATCCAGTCACCTCTGAACATGAA C-3’ | | | 5’-GGAAGATGAATGTCAGAGCTAC AC-3’ | |
| IL-6 | 5’-GACAGCCACTCACCTCTTCA-3’ | | | 5’-CCTCTTTGCTGCTTTCACAC-3’ | |
| IL-10 | 5’-TCAAGGCGCATGTGAACTCC-3’ | | | 5'-GATGTCAAACTCACTCATGGCT- 3’ | |
| JAK1 | 5’-ATTGAGAACGAGTGTCTAGGGA- 3’ | | | 5’-CCTTCAGGTCATGCGTGGAC-3’ | |
| JAK2 | 5’-TCTGGGGAGTATGTTGCAGAA-3’ | | | 5’-AGACATGGTTGGGTGGATACC-3’ | |
| STAT3 | 5’-ATCACGCCTTCTACAGACTGC-3’ | | | 5’-CATCCTGGAGATTCTCTACCACT -3’ | |
| GAPDH | 5’-GCGGGGCTCTCCAGAACATCAT- 3’ | | | 5’-CCAGCCCCAGCGTCAAAGGTG-3’ | |
| **Supplemental Table 2. Cytokine expression levels in human ventricular myocytes (AC16) pro-post irradiation** | | | | | |
| Cytokine | | Control | Irradiation | | *P* value |
| IL-6(pg/ml) | | 2.263±0.05544 | 8.346±0.07740 | | *p*<0.0001 |
| MCP-1(pg/ml) | | 50.28±1.357 | 229.3±0.9208 | | *p*<0.0001 |
| CCL5(ng/ml) | | 4.507±0.2218 | 14.82±0.05199 | | *p*<0.0001 |
| CXCL10(pg/ml) | | 10.96±0.1870 | 60.52±1.031 | | *p*<0.0001 |
| IL-10(pg/ml) | | 159.4±4.165 | 104.0±0.8725 | | *p*<0.0001 |

**Supplemental Table 3. Cytokine expression levels in human ventricular myocytes (AC16) before and after PD-L1 knockdown by siRNA in non-irradiated group**

| Cytokine | IR-siNC | IR-siPD-L1 | *P* value |
| --- | --- | --- | --- |
| IL-6(pg/ml) | 0.9273±0.04666 | 13.31±0.3548 | *p*<0.0001 |
| CCL5(ng/ml) | 5. 114±0.1208 | 22.55±0.6851 | *p*<0.0001 |
| CXCL10(pg/ml ) | 10.46±0.2718 | 50.07±0.3207 | *p*<0.0001 |

**Supplemental Table 4. Cytokine expression levels in human ventricular myocytes (AC16) before and after PD-L1 knockdown by siRNA in irradiated group**

| Cytokine | IR+siNC | IR+siPD-L1 | *P* value |
| --- | --- | --- | --- |
| IL-6(pg/ml) | 0.87±0.06749 | 6.74±0.1107 | *p*<0.0001 |
| CCL5(ng/ml) | 6.74 ± 0.2214 | 17.67±0.5292 | *p*<0.0001 |
| CXCL10(pg/ml | 12.40±0.4618 | 20. 10±0.8406 | *p*<0.0001 |

**Supplemental Fig.1 siRNA **Transfection of AC16****

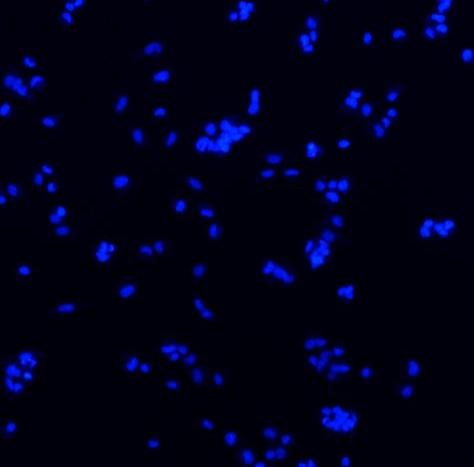

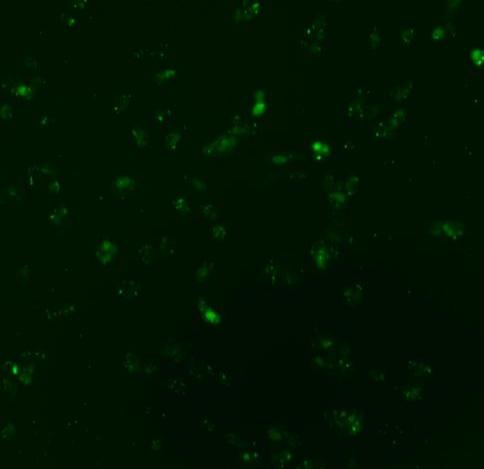

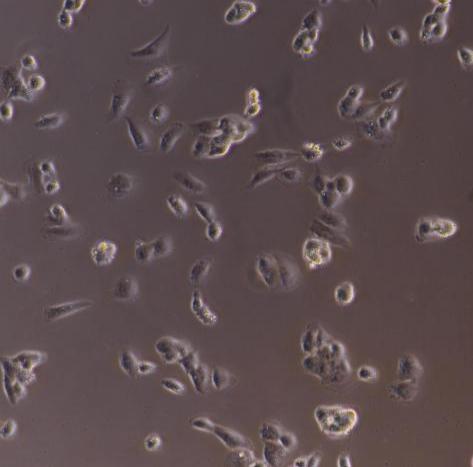

SiRNA	AC16 cells	DAPI

**Supplemental Fig.2**

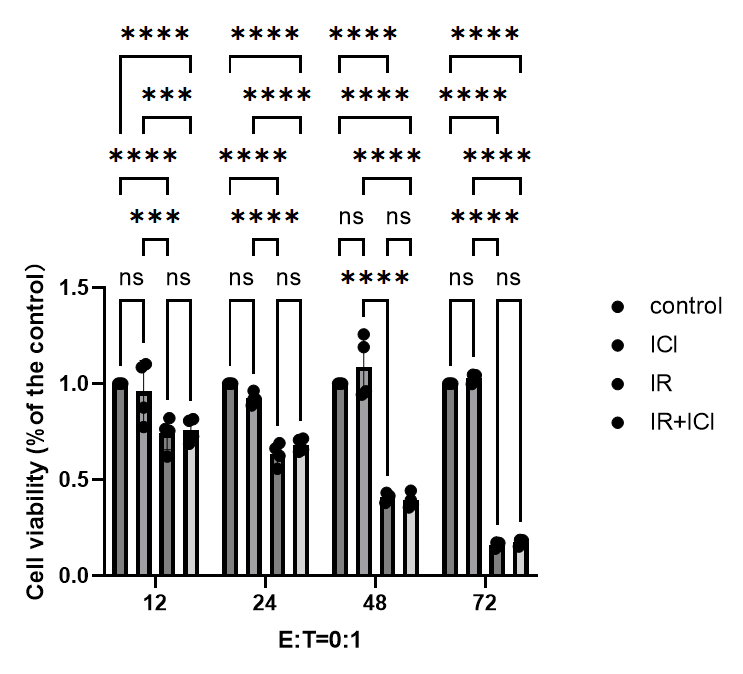

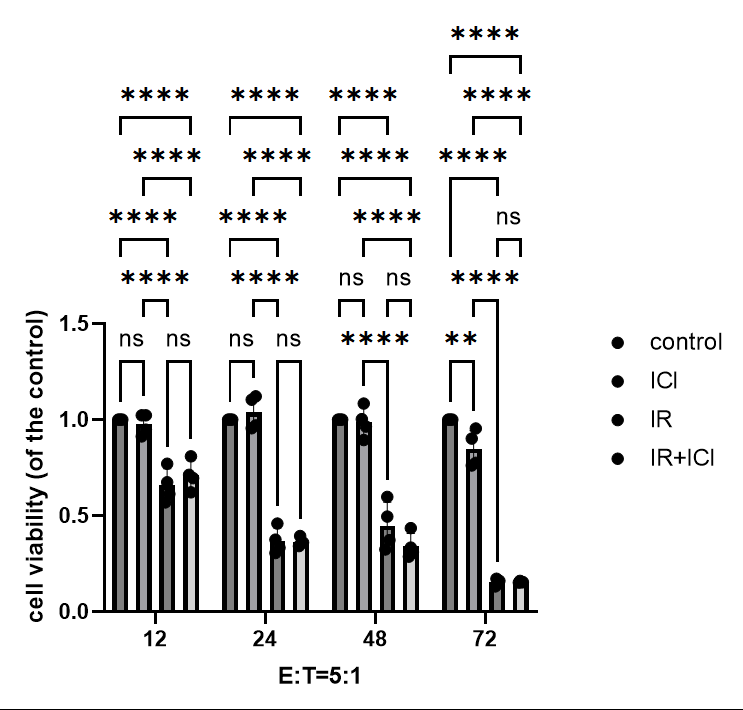

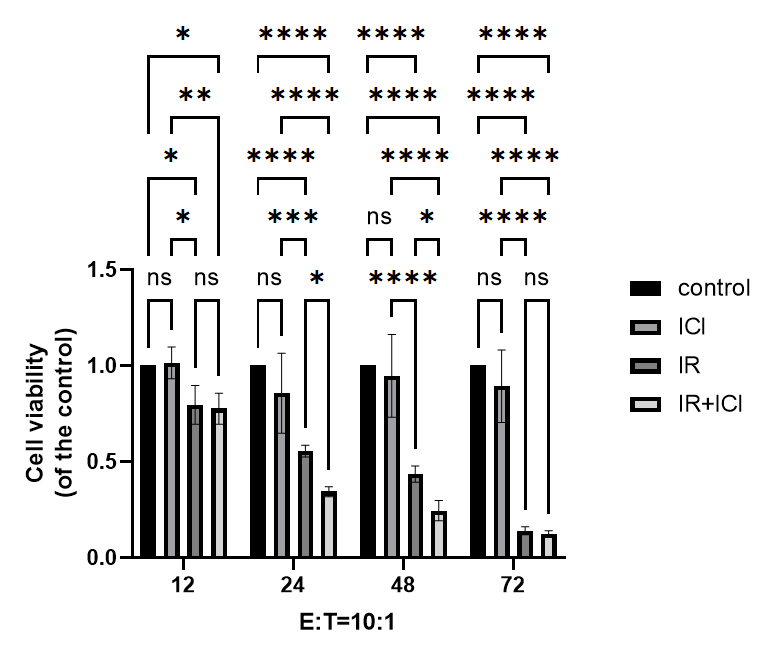

**Supplemental Fig.3 The imaging under microscopic:** **post-irradiation AC16 cells co-cultured with activated CD8+T cells under E:T=0:1,5:1,10:1 and ICI concentrations=5μg/ml for 48h**

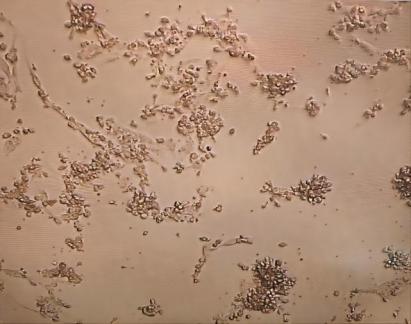

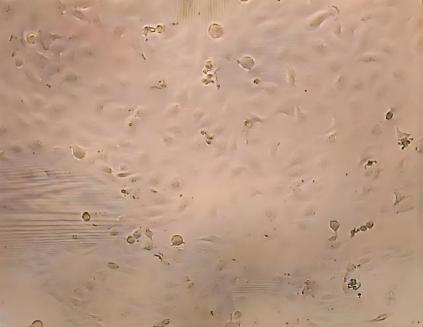

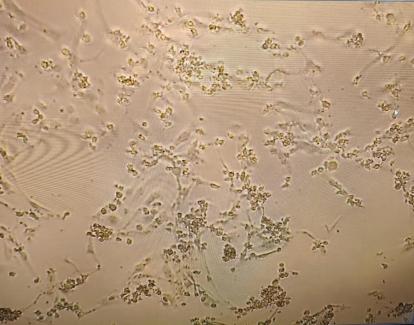

E:T=0:1	E:T=5:1	E:T=10:1

**Supplemental Fig.4 PD-1 expression on CD8⁺ T cells**

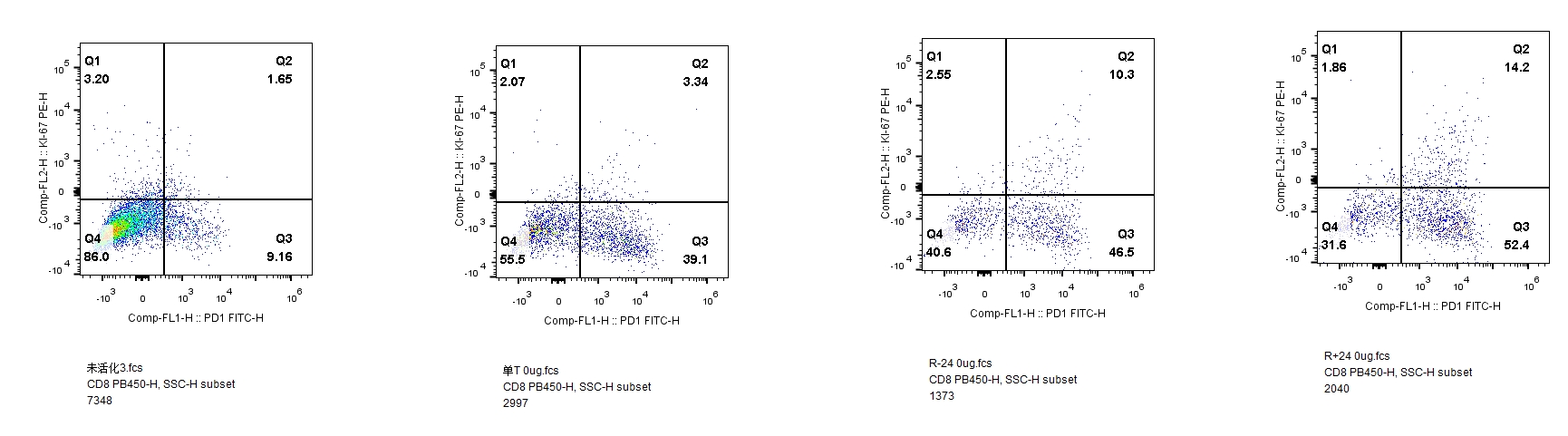

Non-activated CD8⁺ T cells alone

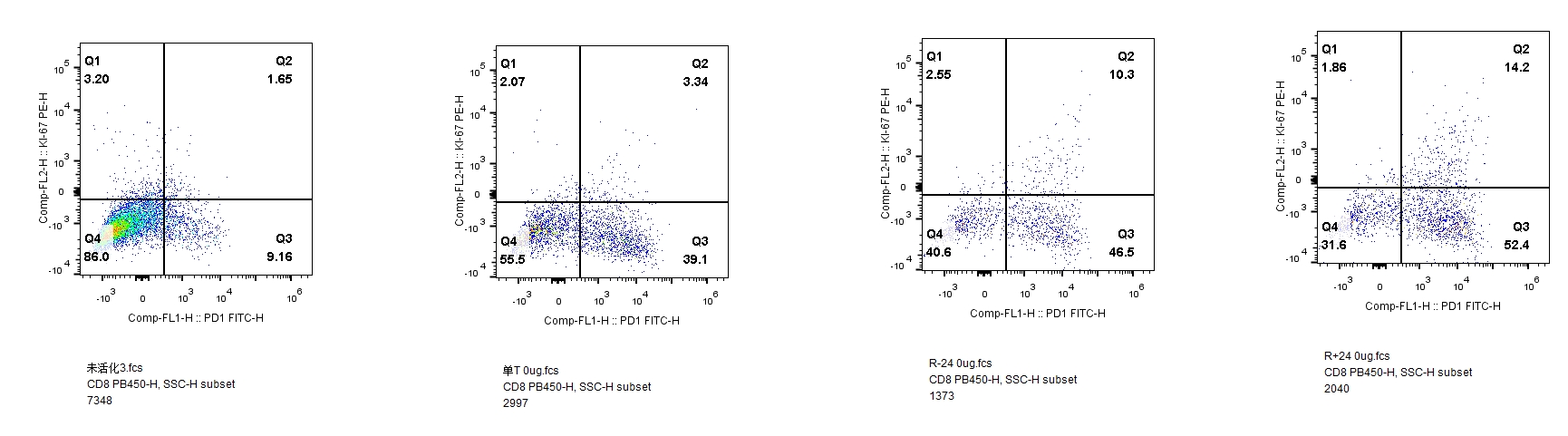

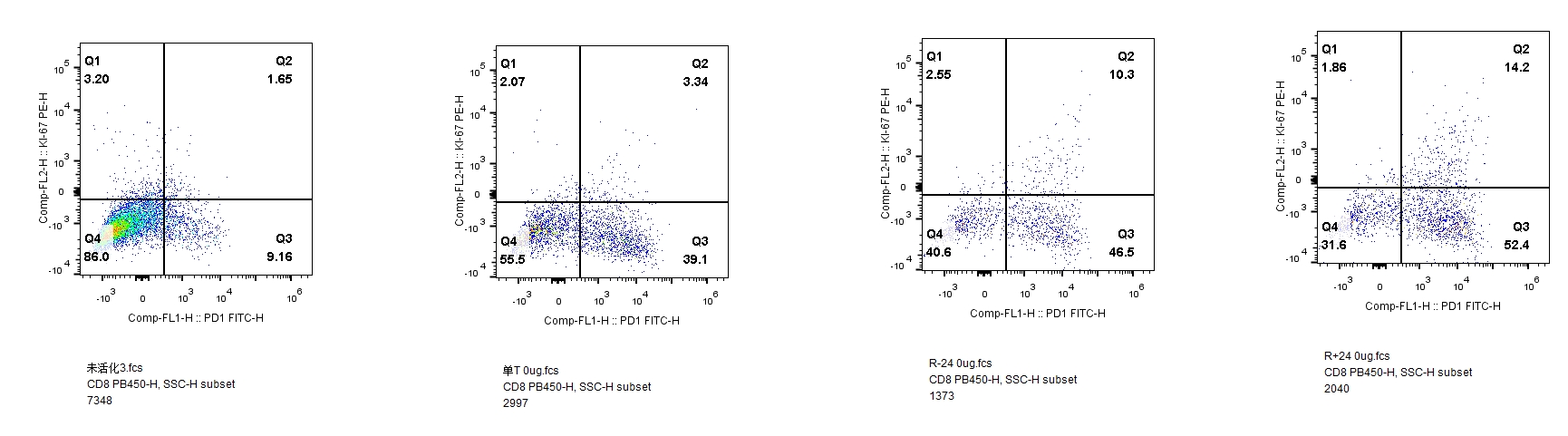

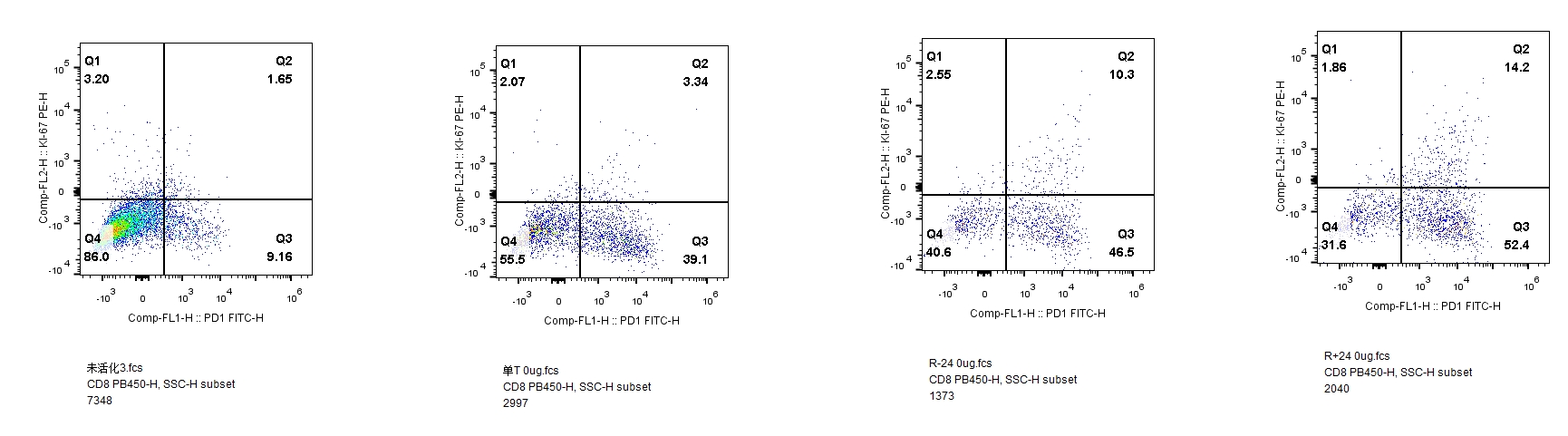

Co-cultured activated CD8⁺ T cells

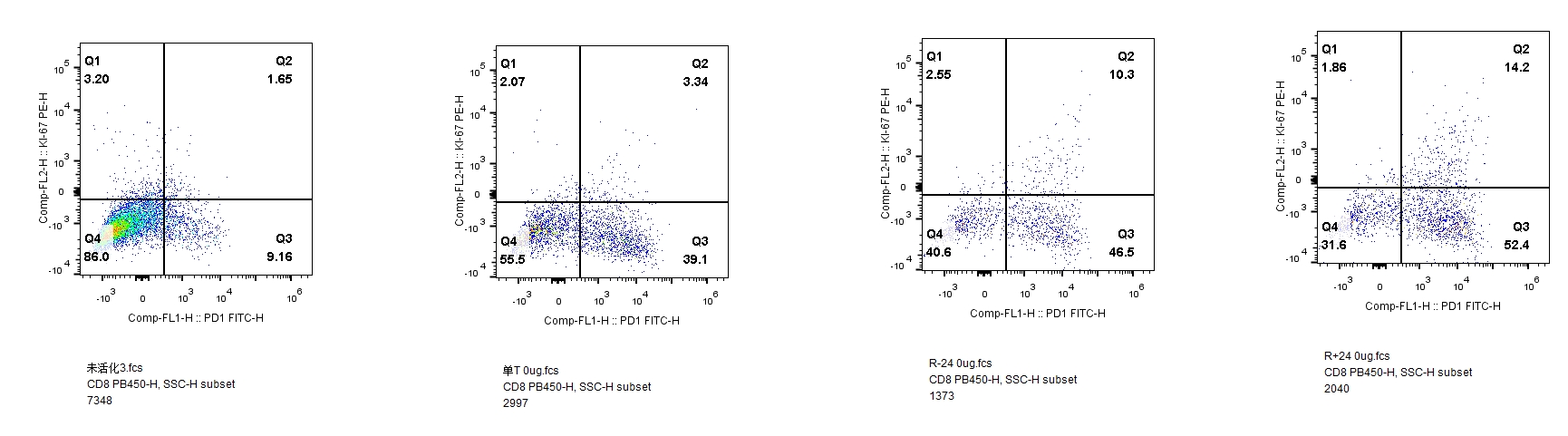
